# VCP inhibition preserves photoreceptor integrity under hydroquinone-induced oxidative stress in a human iPSC-RPE/porcine neuroretina co-culture model

**DOI:** 10.64898/2026.08.18.745423

**Authors:** Ana-Cristina Almansa-García, Angela Armento, Shibu Antony, Mohamed-Ali Jarboui, Rosario Fernández-Godino, Enrico Cossio, Bowen Cao, Anne-Sophie Petremann-Dumé, Anneli Vollert, Ellen Kilger, Sylvie Bolz, Marius Ueffing, Blanca Arango-Gonzalez

## Abstract

Age-related macular degeneration (AMD) is the leading cause of irreversible vision loss in older adults. It is characterised by early retinal pigment epithelium (RPE) dysfunction followed by progressive photoreceptor degeneration. Cigarette smoking is a major environmental risk factor for AMD, and hydroquinone (HQ), a redox-active cigarette smoke component, induces oxidative stress and apoptosis in RPE cells. To analyse how RPE stress contributes to photoreceptor degeneration, we employed a retinal co-culture model composed of human induced pluripotent stem cell-derived RPE (iPSC-RPE) cells in conjunction with porcine neuroretina explants. Exposure to HQ induced oxidative stress in iPSC-RPE cells as well as retinal photoreceptors (RPR), resulting in apoptosis, executed at least in part by caspase activation. Concomitantly, HQ caused endoplasmic reticulum (ER) stress (ERAD) in RPR followed by their degeneration, evidenced by reduced outer nuclear layer (ONL) rows and shortened RPR outer segments (OS). Based on earlier results, which suggest a perturbation of proteostasis due to HQ, we tested whether ML240, a bona fide inhibitor of valosin-containing protein (VCP), would influence the degree of degenerative activities. ML240 did not prevent HQ-induced apoptosis in iPSC-RPE cells. However, it significantly preserved photoreceptor integrity, retaining OS length and cone density in HQ-stressed co-cultures.

Proteomic analysis suggested that ML240 reshapes stress response patterns of the HQ-exposed neuroretina, as evidenced by a reduction in ERAD-associated markers, increased levels of antioxidant response proteins, and the preservation of cytochrome c enrichment in photoreceptor inner segments, which indicates improved mitochondrial integrity consistent with the observed preservation of photoreceptor structure.

Together, these findings establish the iPSC-RPE/neuroretina co-culture as a platform to analyse pathophysiological features of AMD, dissect cell type-specific retinal responses to environmental stress and test neuroprotective pharmacological approaches to protect photoreceptors in oxidative stress-associated retinal degeneration.

## Introduction

Age-related macular degeneration (AMD) is the leading cause of irreversible vision loss in older adults, affecting approximately 200 million people worldwide with projections approaching 300 million by 2040 [1, 2]. Early AMD is characterized by drusen, extracellular lipid- and protein-rich deposits located between the retinal pigment epithelium (RPE) and Bruch’s membrane, along with functional and structural abnormalities of the RPE. Advanced AMD manifests as neovascular AMD, characterized by choroidal neovascularization, or as geographic atrophy, characterized by progressive loss of the RPE and photoreceptors [3]. AMD primarily affects the macula, the cone-enriched central retina responsible for high-acuity vision. As the disease progresses, loss of central visual acuity compromises daily activities such as reading, facial recognition, or driving [3].

AMD is a multifactorial disease caused by aging, genetic susceptibility, environmental exposures, and lifestyle factors, and involves multiple retinal, immune and choroidal cell types [4]. Because RPE dysfunction and alterations in the sub-RPE microenvironment are early features of AMD, many experimental studies have focused primarily on RPE-centered mechanisms. Although AMD pathogenesis remains incompletely understood, complement dysregulation, mitochondrial dysfunction, metabolic imbalance, impaired antioxidant defense, defective proteostasis, and inflammatory signalling have been implicated in RPE pathology [5–7]. By contrast, neuroretinal responses to RPE stress remain less investigated, despite the fact that photoreceptor degeneration ultimately drives irreversible visual dysfunction. This gap is partly due to the limited availability of human-relevant models that preserve RPE-neuroretina interactions.

Cigarette smoking is one of the strongest modifiable environmental risk factors for AMD [8]. Hydroquinone (HQ), a redox-active benzene metabolite and cigarette smoke component, promotes the generation of reactive oxygen species (ROS), oxidative stress, and DNA damage [9]. HQ has therefore been widely used across various *in vitro* and *in vivo* systems to model cigarette smoke-related injury in AMD, mainly in the context of drusen-like deposit formation and RPE pathology [10–13]. However, it is not established whether HQ is sufficient to drive photoreceptor damage in a RPE-neuroretina co-culture model, which retains complex tissue-like cell interactions.

In response to oxidative stress, other cellular compartments functions are impaired such as the endoplasmic reticulum (ER), involved in regulating protein folding, calcium homeostasis, and lipid biosynthesis [14, 15]. Because protein folding depends on a tightly regulated ER redox environment, oxidative stress can impair folding capacity and perturb proteasomal degradation pathways [16]. ER stress is considered to play an important role in retinal degenerative disorders, including AMD, through its close association with oxidative stress, proteostasis defects, inflammation, and aging [17, 18]. Thus, modulation of ER stress response and the proteasome activity may represent a therapeutic strategy to counteract oxidative stress-induced retinal damage [19, 20].

Valosin-containing protein (VCP/p97), an ATP-dependent protein remodeling factor involved in proteostasis, ER-associated degradation, and mitochondrial quality control, has emerged as a potential therapeutic target in retinal degenerative disorders characterized by defects in proteostasis and energy homeostasis [21–23]. However, whether VCP inhibition can protect photoreceptors under HQ-driven oxidative stress remains unknown.

In this study, we used HQ as an AMD-relevant environmental stressor in a human iPSC-RPE/porcine neuroretina co-culture model [24]. HQ was applied alone or in combination with the VCP inhibitor ML240 [24] to determine whether VCP inhibition modulates RPE stress responses and preserves photoreceptor integrity in the co-cultured neuroretina.

## Results

### Establishment of a human iPSC-RPE/porcine neuroretina co-culture model for oxidative stress studies

We employed an AMD-relevant co-culture model that combines human induced pluripotent stem cell-derived retinal pigment epithelium (iPSC-RPE) with porcine neuroretinal explants, with the photoreceptors facing the RPE cells, thereby preserving direct interactions between the RPE and photoreceptor outer segments (**Figure 1A**) [25]. Human iPSCs were differentiated into RPE cells using a previously established protocol [13] and matured until they displayed characteristic RPE features, including pigmentation, cobblestone morphology, functional trans-epithelial electrical resistance (TER), and polarized secretion of pigment epithelium-derived factor (PEDF) (**Figure 1B-E**). Porcine neuroretina explants were isolated from the cone-rich visual streak, a region that partially resembles the photoreceptor composition of the human parafovea (**Figure 1A**).

**Figure 1.**
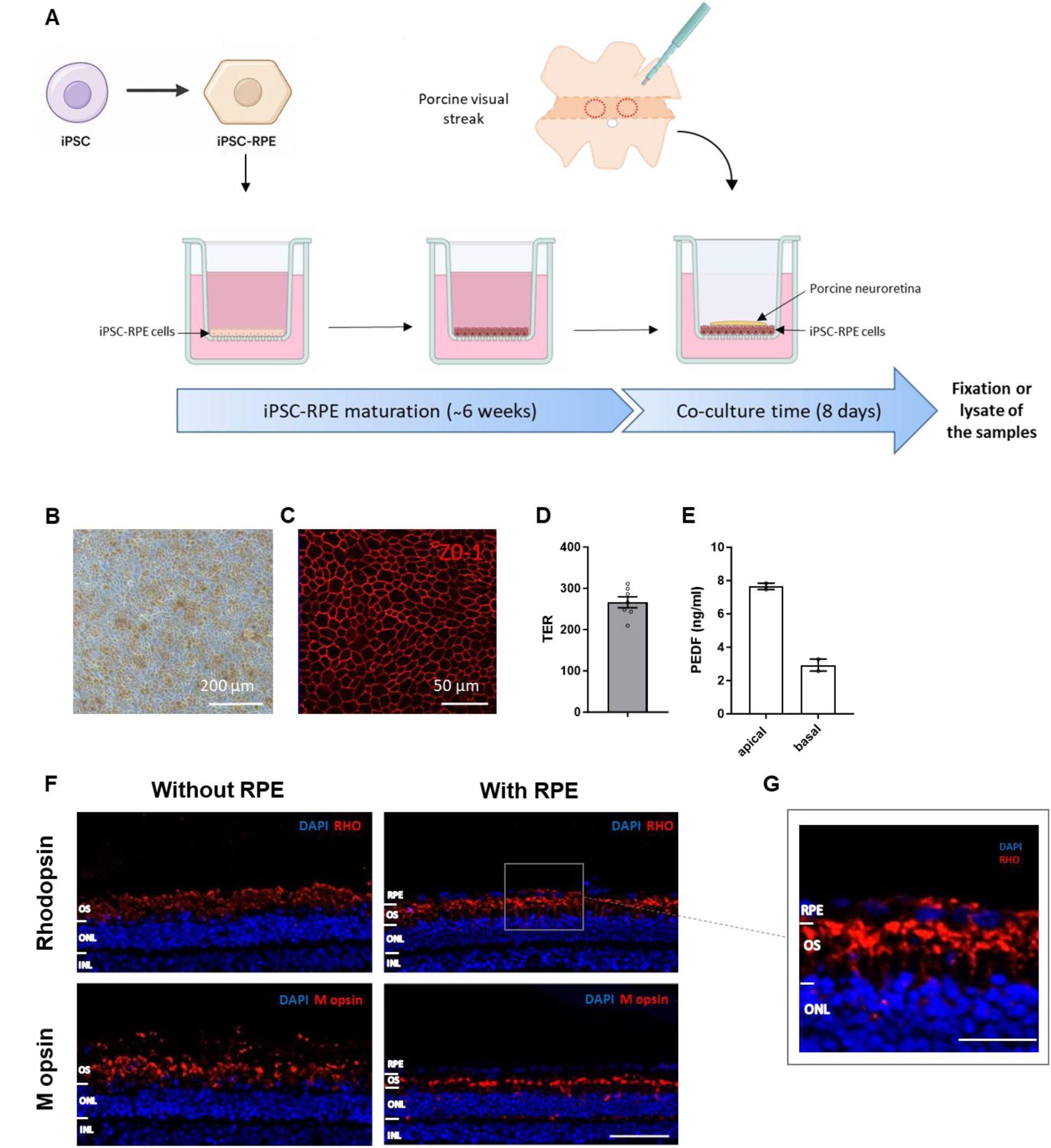
Characterization of the human iPSC-RPE/porcine neuroretina co-culture system. (**A**) Schematic representation of the co-culture setup and experimental workflow. (**B**) Light microscopy image of differentiated iPSC-RPE cells showing characteristic pigmentation and cobblestone morphology. (**C**) Immunofluorescence staining of the tight junction marker ZO-1. (**D**) Transepithelial electrical resistance (TER) measurements of mature iPSC-RPE monolayers. (**E**) Quantification of apical and basal secretion of pigment epithelium-derived factor (PEDF) by ELISA. (**F**) Representative fluorescence images of porcine neuroretinal explants cultured for 8 days with or without iPSC-RPE cells and stained for rhodopsin (rods) or M opsin (cones), and counterstained with DAPI. (**G**) High-magnification image showing rhodopsin immunostaining at the iPSC-RPE/neuroretina interface.

Porcine retinal explants were cultured in vitro for 8 days with or without co-cultured iPSC-RPE cell. Both groups showed typical retinal structure and organization, however, co-cultured explants showed better preserved photoreceptor outer segment (OS) organization than explants cultured without iPSC-RPE cells (**Figure 1F**). Rhodopsin immunostaining at the RPE/OS interface supported physical contact and interaction between iPSC-RPE cells and retinal explants (**Figure 1G**).

To model exposure to an AMD-relevant environmental stressor, co-cultures were treated with HQ. First, iPSC-RPE monocultures were treated for 48 h with 100 µM HQ or 0.4% DMSO as a vehicle control. This HQ concentration was selected based on previous *in vitro* studies in iPSC-RPE cells [13]. We confirmed that the iPSC-RPE line used in this study was susceptible to HQ-induced stress, as shown by elevated caspase activity and oxidative stress (**Figure 2A-B**). Western blot analysis further showed an increased ratio of the proapoptotic protein BAX to the anti-apoptotic protein BCL2 in HQ-treated iPSC-RPE cells compared with controls (**Figure 2C-D**). Cleaved caspase-3 immunofluorescence after 8 days of treatment further supported sustained apoptotic activation in HQ-exposed iPSC-RPE cells (**Figure 2E**). Together, these data indicate that HQ shifts iPSC-RPE cells toward apoptosis.

**Figure 2.**
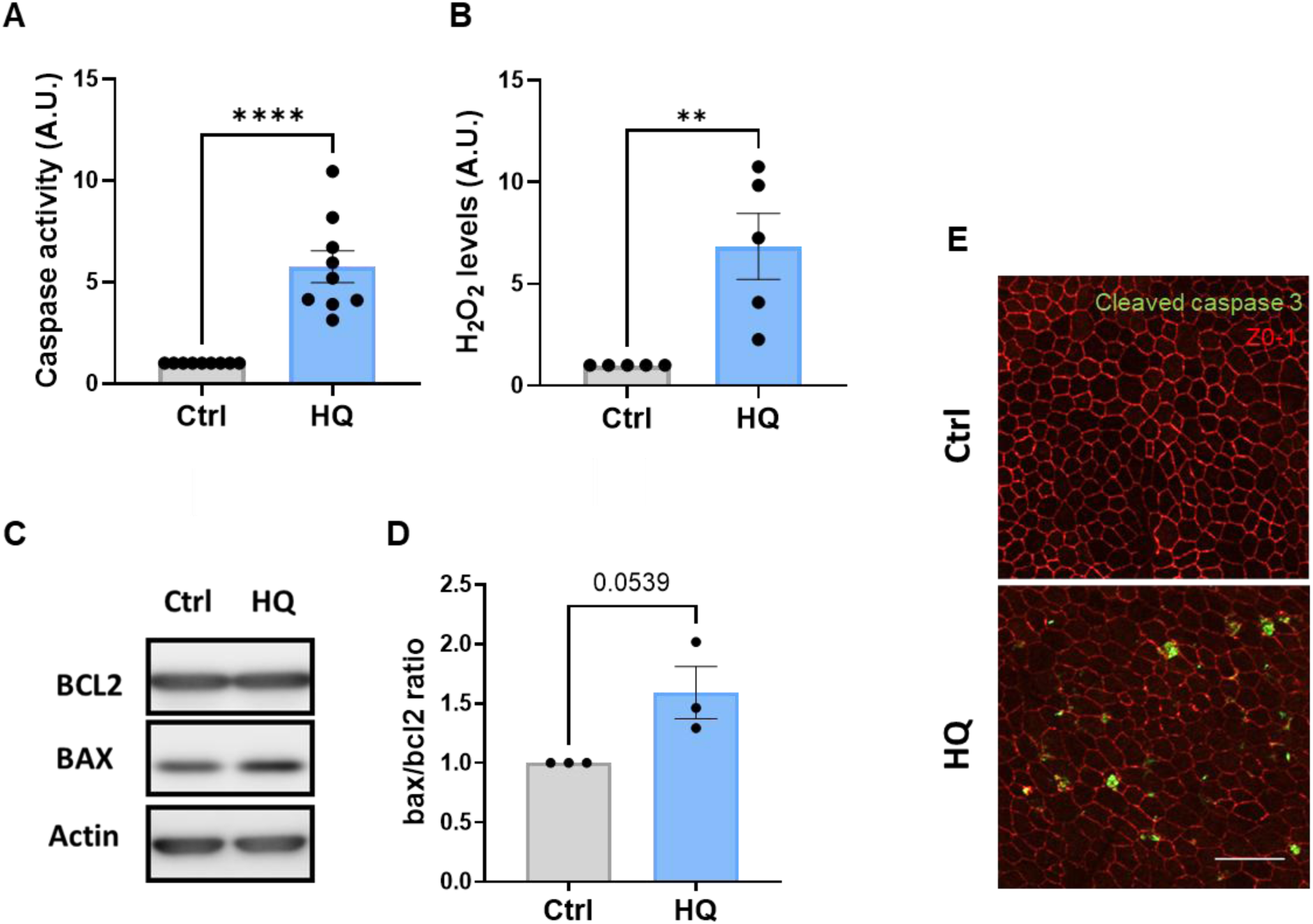
Hydroquinone (HQ) induces oxidative stress and apoptotic activation in iPSC-RPE cells. (**A**) Caspase-3/7 activity, (**B**) H_2_O_2_ levels, as a measure of oxidative stress, were assessed after 48 h of treatment with 100 µM HQ or 0.4% DMSO control. (**C**) Representative Western blot images showing BCL2, BAX, and β-actin (loading control). (**D**) Quantification of the BAX/BCL2 ratio. (**E**) Immunofluorescence staining for cleaved caspase-3 (green) and ZO-1 (red) in iPSC-RPE cells after 8 days of HQ treatment. Differences between groups were determined using unpaired Student’s t-test (n = 6-9 biological replicates for caspase and oxidative stress assays; n = 3 for Western Blot). Data are presented as mean ± SEM. ** p ˂ 0.01. **** p ˂ 0.0001.

### Hydroquinone induces photoreceptor degeneration in the human iPSC-RPE/porcine retina co-culture model

We next investigated whether HQ was associated with photoreceptor damage in the complete human iPSC-RPE/porcine neuroretina co-culture. Co-cultures were treated with HQ for 8 days *in vitro* (DIV 8). Photoreceptor integrity was assessed by outer nuclear layer (ONL) row counts, photoreceptor outer segment (OS) length, and cone density measurements. HQ-treated co-cultures showed significantly fewer ONL rows (**Figure 3A, C**), reduced OS length (**Figure 3A, D**), and a non-significant decrease in cone photoreceptor density (**Figure 3B, E**), compared with vehicle-treated controls. These data show that, in the co-culture system, exposure to HQ is associated not only with RPE damage, but also with photoreceptor degeneration in the co-cultured porcine neuroretina. Together with the iPSC-RPE data, these findings indicate that HQ exposure induces a combined RPE and photoreceptor degenerative phenotype in the co-culture system.

**Figure 3.**
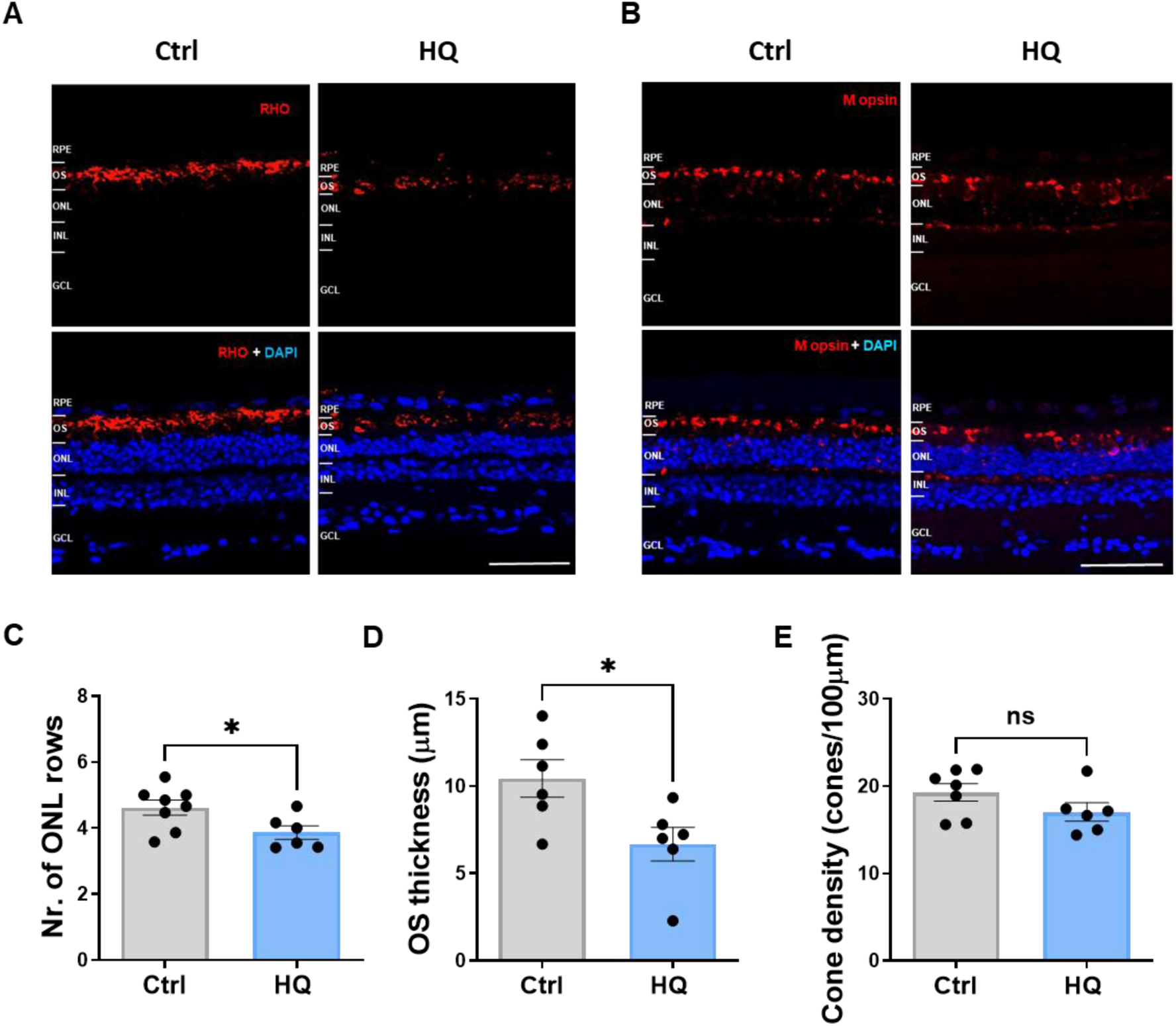
HQ induces photoreceptor degeneration in the human iPSC-RPE/porcine retina co-culture model. Porcine neuroretinal explants were co-cultured with mature iPSC-RPE cells for 8 days (DIV 8) and treated with 100 µM HQ or 0.4% DMSO (vehicle control). Representative images show rod and cone photoreceptors visualized with (**A**) rhodopsin and (**B**) M opsin, respectively, with DAPI counterstaining. (**C**) Quantification of outer nuclear layer (ONL) cell rows. (**D**) Photoreceptor outer segment (OS) layer thickness measurements. (**E**) Cone density quantification. Differences between HQ and control groups were analyzed using unpaired Student’s t-test (n = 6-8 biological replicates). Data presented as mean ± SEM. * p ˂ 0.05. ns = not significant.

### VCP inhibition selectively protects photoreceptors from HQ-induced damage

Treatment with the cigarette smoke component HQ showed induced stress in iPSC-RPE cells and photoreceptor degeneration in the co-cultured neuroretinas. This system thus models environmental stress-related retinal damage as also observed in AMD and can serve as a platform to test the effect of potential therapeutic interventions. We next evaluated the effect of VCP inhibition, an emerging neuroprotective approach in retinal degenerative diseases. Using the co-culture system, we tested whether pharmacological inhibition of VCP with ML240 could attenuate the HQ induced retinal damage. First, iPSC-RPE monocultures were treated for 48 h with 100 µM HQ alone or in combination with 20 µM ML240. VCP inhibition by ML240 did not significantly reduce HQ-induced caspase activity or oxidative stress in iPSC-RPE cells after 48 h of treatment (**Figure 4A-B**). Consistently, cleaved caspase-3 immunofluorescence after 8 days of treatment showed comparable apoptotic activation in HQ-and HQ plus ML240-treated iPSC-RPE cells (**Figure 4C**). Thus, ML240 treatment did not prevent HQ-induced damage in iPSC-RPE cells.

**Figure 4.**
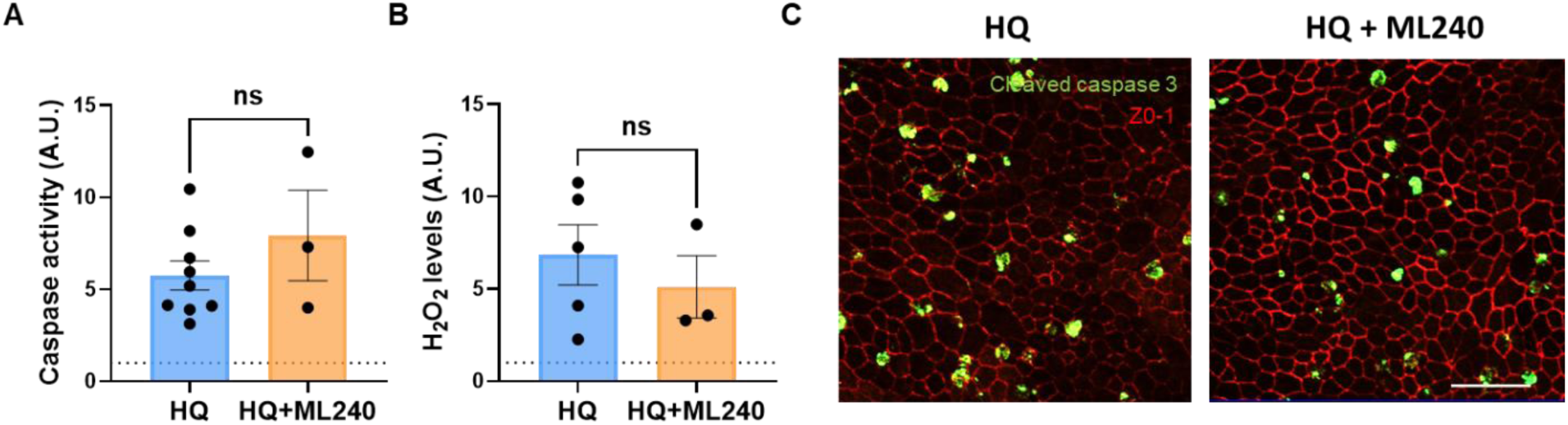
Effect of VCP inhibition by ML240 in HQ-stressed iPSC-RPE cells. (**A**) Caspase-3/7 activity, and (**B**) H_2_O_2_ levels were quantified after 48 h of treatment with 100 µM HQ or 100 µM HQ + 20 µM ML240. (**C**) Immunofluorescence staining for cleaved caspase-3 (green) in iPSC-RPE cells after 8 days of treatment with 100 µM HQ or 100 µM HQ + 20 µM ML240. Differences between HQ and HQ + ML240 groups were assessed using an unpaired Student’s t-test (n = 3-8 biological replicates). Data presented as mean ± SEM; ns = no significance.

Next, we evaluated whether ML240 could prevent photoreceptor degeneration in HQ-stressed co-cultured neuroretinas. Photoreceptor integrity was assessed by rhodopsin and M opsin immunofluorescence to visualize rods and cones, respectively (**Figure 5A–B**). Compared with HQ-treated co-cultures, HQ plus ML240-treated co-cultures showed significantly increased OS length (**Figure 5D**) and cone density (**Figure 5E**), with a non-significant trend toward increased ONL cell rows (**Figure 5C**). Together, these results indicate that ML240 does not prevent HQ-induced apoptotic activation in iPSC-RPE cells but selectively preserves photoreceptor structure in HQ-stressed co-cultured neuroretinas.

**Figure 5.**
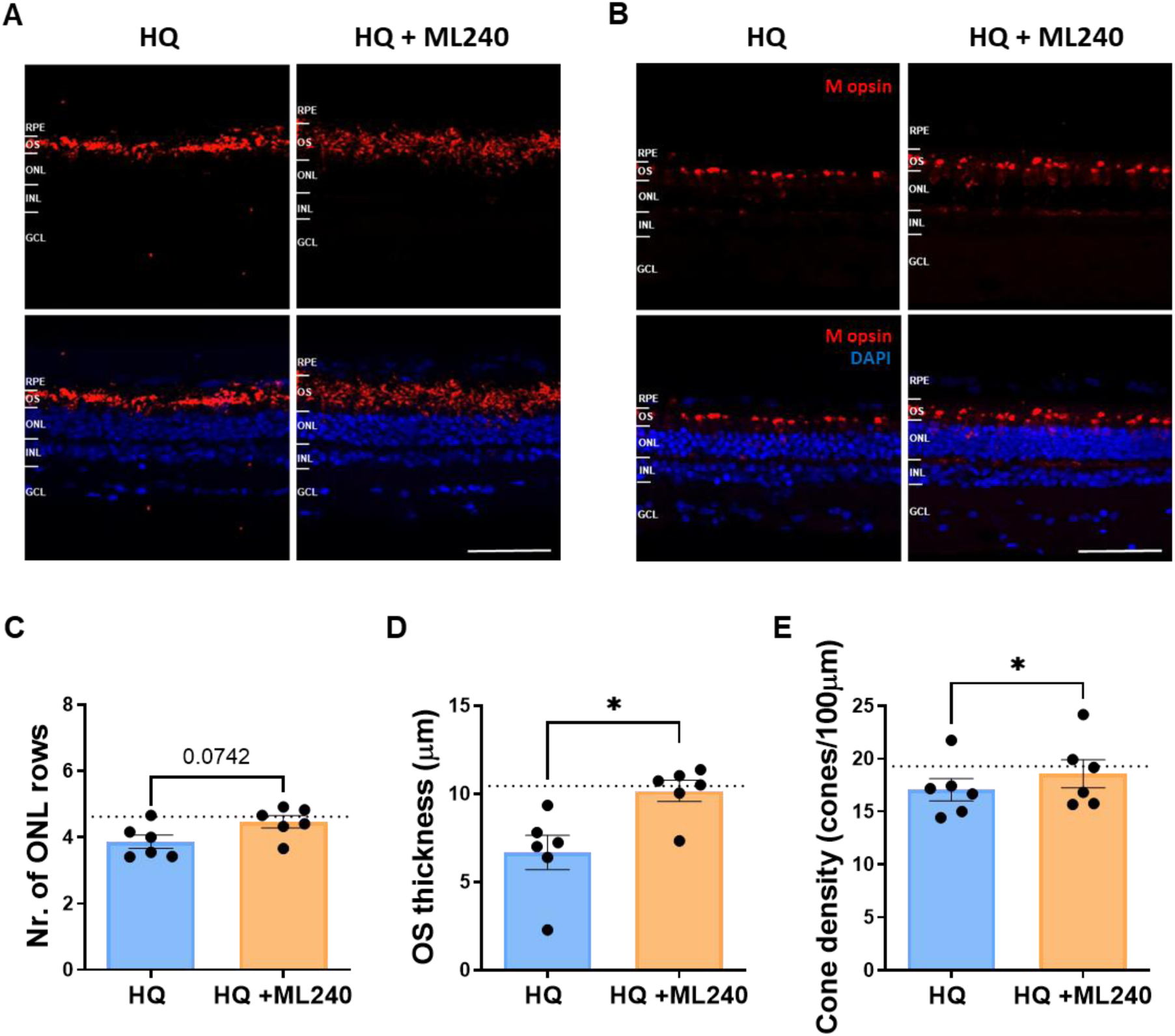
VCP inhibition preserves photoreceptor integrity in HQ-stressed human iPSC-RPE/porcine retina co-culture model. Porcine neuroretina explants were co-cultured with mature iPSC-RPE cells for 8 days (DIV 8) and treated with 100 µM HQ alone or with 100 µM HQ + 20 µM ML240. (**A**) Rhodopsin (red) and (**B**) M opsin (red) immunostaining visualized rod and cone photoreceptors, respectively, and DAPI (blue) was used as counterstaining. (**C**) Quantification of ONL cell rows. (**D**) Measurement of photoreceptor OS layer thickness. (**E**) Quantification of cone density. Differences between the HQ and HQ + ML240 groups were assessed using a paired Student’s t-test (n = 6 biological replicates). Data presented as mean ± SEM. * p ˂ 0.05.

### ML240 reshapes the proteomic stress response of HQ-exposed co-cultured retinas

Because ML240 preserved photoreceptor structure in HQ-stressed co-cultures without attenuating apoptotic activation in iPSC-RPE cells, we next investigated neuroretina-specific molecular responses. Porcine retina/iPSC-RPE co-cultures were treated with HQ alone or HQ + ML240 for 8 days. Neuroretinas were then fixed for immunohistochemistry or separated from the RPE and processed for mass spectrometry or Western blot analysis.

Mass spectrometry revealed differential protein abundance between HQ-stressed neuroretinas treated with or without ML240 (**Figure 6A, Supplementary table 1**). Performing GO enrichment analysis, pathway enrichment analysis showed that proteins increased in both groups were primarily associated with cellular responses to stress or stimuli, but the specific proteins and pathways differed between HQ and HQ + ML240 treatment groups (**Figure 6B, C, Supplementary table 2**).

**Figure 6.**
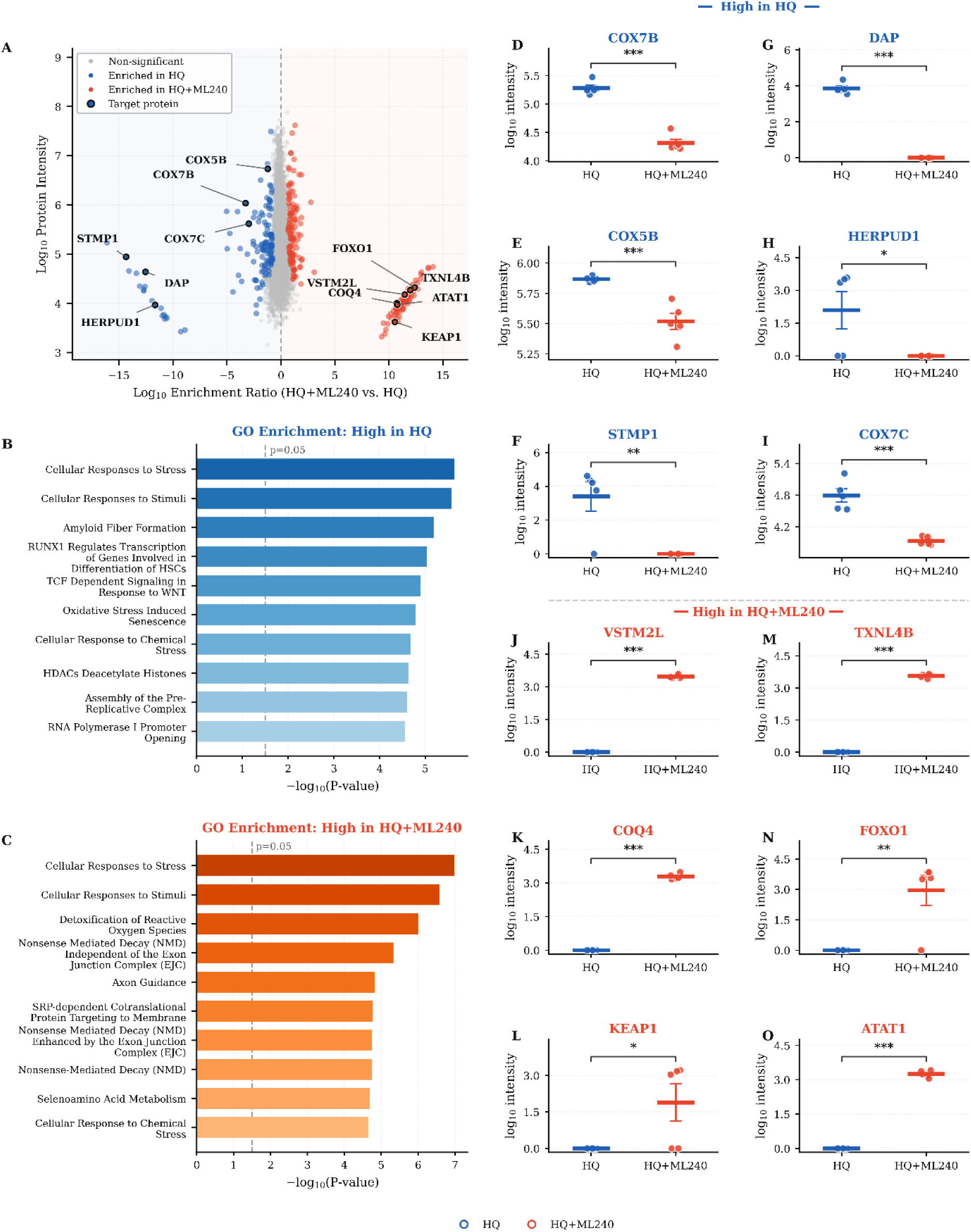
Proteomic remodeling of HQ-stressed co-cultured retinas after ML240 treatment. Porcine neuroretina explants were co-cultured with mature iPSC-RPE cells for 8 days (DIV 8) and treated with 100 µM HQ or HQ + 20 µM ML240. At the end of the co-culture period, neuroretinas were isolated from the iPSC-RPE cells and lysed for mass spectrometry analysis. (**A**) Scatter plot showing the differential protein abundance between HQ-stressed retinal explants and the retinal explants treated with HQ+ML240. Differences are expressed as log10 abundance ratios between HQ+ML240 and HQ treatment groups on the x-axis, with protein abundance shown on the y-axis. Not significantly changed proteins are represented in grey. (**B**) Bar plots showing the top 10 significantly enriched Reactome pathways proteins increased in (B) HQ-treated retinas and (C) HQ+ML240-treated retinas, using the Reactome Pathways 2024 dataset. (D-O) Box plots showing differential abundance of selected proteins involved in enriched pathways between HQ and HQ+ML240 treatment groups. Data are presented as mean ± SEM. * p ˂ 0.05. ** p ˂ 0.01. *** p ˂ 0.001.

Proteins increased in the HQ-treated neuroretinas relative to HQ+ML240 were associated with apoptotic processes, including death-associated protein (DAP, **Figure 6G**), and with mitochondrial respiratory chain components, including cytochrome c oxidase subunits COX7B, COX5B and COX7C, as well as short transmembrane mitochondrial protein 1 (STMP1; Figure **6D-F**, **I**). This profile suggests a stress-associated mitochondrial remodeling response in HQ-exposed neuroretinas.

In addition, HQ-treated explants showed increased abundance of proteins linked to ER-associated degradation (ERAD) and the ubiquitin-proteasome system, including homocysteine inducible ER protein with ubiquitin-like domain 1 (HERPUD1, **Figure 6H**), heat shock protein 90 beta family member 1 (HSP90B1), proteasome 20S subunit beta 6 (PSMB6), and ubiquitin (UBC) (**Supplementary table 2**). These changes are consistent with increased proteostatic stress and activation of protein quality-control pathways in HQ-stressed neuroretinas compared with HQ + ML240-treated neuroretinas.

On the other hand, HQ+ML240-treated neuroretinas showed increased abundance of proteins associated with mitochondrial homeostasis, antioxidant defence, and cytoskeletal or vesicle-trafficking pathways. These included the voltage-dependent anion channel 1 (VDAC1), binding protein V-set and transmembrane domain-containing 2-like (VSTM2L) and coenzyme Q4 (COQ4; **Figure 6J, K**) as well as antioxidant and redox-associated proteins such as thioredoxin-like 4B (TXNL4B), forkhead box O1 (FOXO1), Kelch-like ECH-associated protein 1 (KEAP1) (**Figure 6L-N**), glutathione S-transferase pi 1 (GSTP1), peroxiredoxin 1 (PRDX1), superoxide dismutase 1 (SOD1) or catalase (CAT) (**Supplementary table 2**). Proteins related to microtubule acetylation and vesicle trafficking, such as alpha tubulin acetyltransferase 1 (ATAT1, **Figure 6O**), IQ motif containing GTPase-activating protein 3 (IQGAP3), kinesin family member 1C (KIF1C), or Ras-related protein Rab-39A (RAB39A), were also increased (**Supplementary table 2**).

These proteomic changes suggest that ML240 redirects the HQ-induced neuroretinal stress response away from ERAD/proteasome-associated stress markers and toward antioxidant, mitochondrial, and cytoskeletal homeostatic programs, consistent with the observed preservation of photoreceptor structure.

### ML240 attenuates HQ-induced cytochrome c mislocalization in photoreceptors

In the proteomic analyses, the HQ+ML240 treatment group showed increased abundance of selected apoptosis-and mitochondria-associated proteins, including Bcl-2 homologous antagonist/killer (BAK) and total cytochrome c (CYCS) (**supplementary table 1, 2**). Because cytochrome c release from mitochondria to the cytosol is a key event in mitochondrial-related apoptosis [26], we next assessed total cytochrome c protein levels and its localization in co-cultured retinal explants. Therefore, we performed Western blot and immunohistochemical analyses to assess total cytochrome c protein levels and its spatial distribution within the co-cultured retinal explants. The mitochondrial outer membrane protein TOM20 was used as a readout of total mitochondrial content.

Western blot analysis showed that total cytochrome c protein levels were significantly elevated in both HQ and HQ + ML240 treated neuroretinas compared with controls (**Figure 7A, B**). However, cytochrome c immunostaining revealed distinct spatial distribution between treatment groups. In HQ-treated retinas, cytochrome c immunofluorescence extended into the ONL instead of remaining enriched in photoreceptor inner segments (IS), where photoreceptor mitochondria are predominantly localized. This abnormal distribution was attenuated by ML240, as reflected by a higher IS/ONL cytochrome c fluorescence ratio in HQ plus ML240-treated retinas compared with HQ-treated retinas (**Figure 7D-E**). TOM20 protein levels remained unchanged across treatment groups (**Figure 7A, C**), suggesting that the altered cytochrome c distribution was not explained by gross changes in mitochondrial content. Altogether, these findings suggest that ML240 attenuates HQ-induced cytochrome c mislocalization and supports mitochondrial integrity in photoreceptors under oxidative stress.

**Figure 7.**
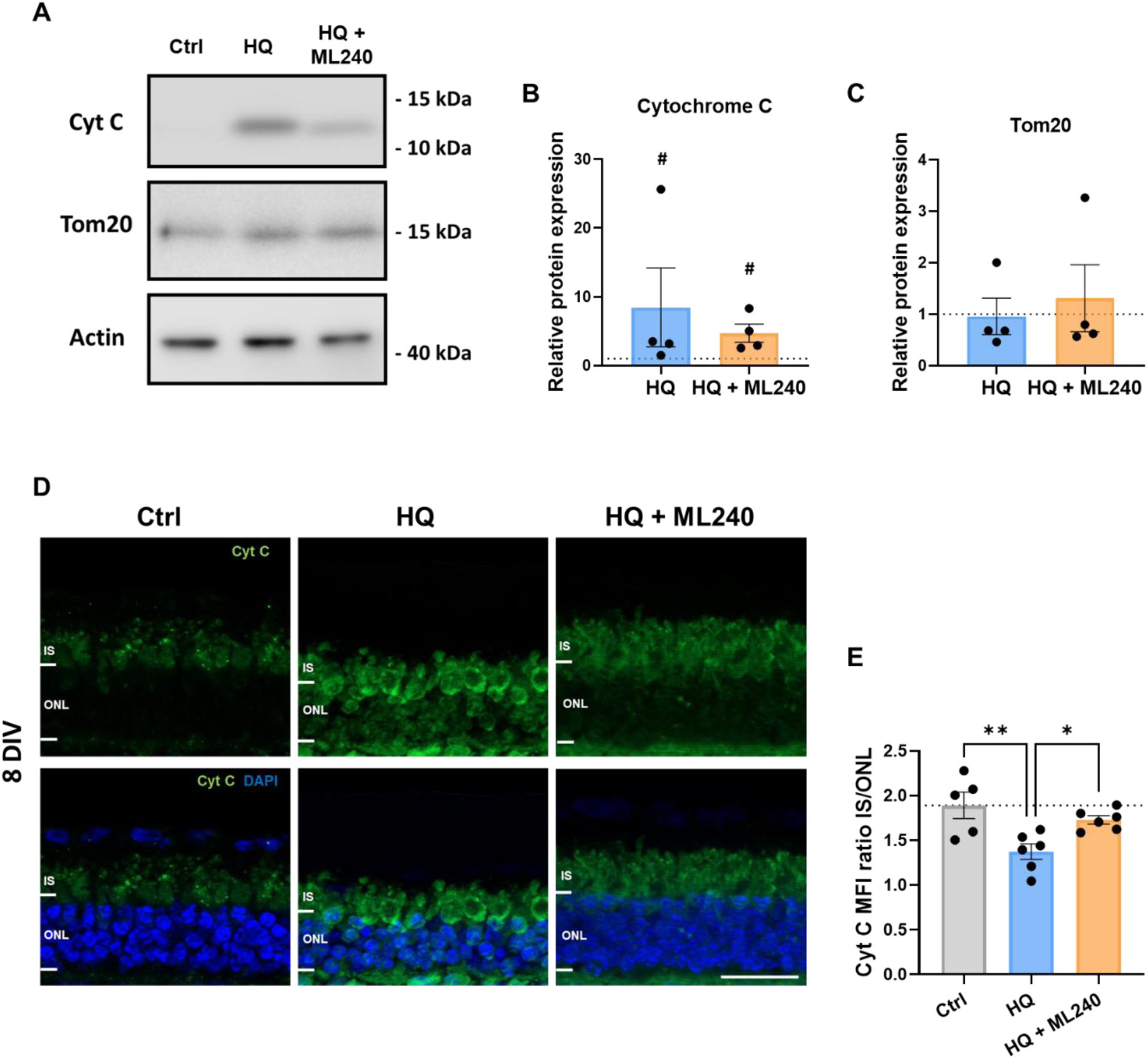
ML240 attenuates HQ-induced cytochrome c mislocalization in co-cultured neuroretinas. Porcine neuroretinal explants were co-cultured with mature iPSC-RPE cells for 8 days (DIV 8) and treated with 100 µM HQ or HQ + 20 µM ML240. (**A**) Representative Western blots and quantification of relative protein levels of (**B**) cytochrome c, and (**C**) TOM20. β-actin was used as a loading control. Differences between treatment groups were assessed by paired or unpaired Student’s t-test according to the experimental design (n = 4 biological replicates). Data presented as mean ± SEM. The control mean is indicated by a dotted line. # p ˂ 0.05 compared to control (dotted line). (**D**) Representative cytochrome c immunofluorescence staining (green) in retinal sections showing altered localization after HQ and HQ + ML240 treatment compared with control. (**E**) Quantification of the mean fluorescence intensity (MFI) ratio between the photoreceptor inner segments (IS) and the ONL. Differences between treatment groups were assessed by one-way ANOVA with Tukey’s multiple comparison test. Data are presented as mean ± SEM. The control average is represented by a dotted line for reference. * p ˂ 0.05. ** p ˂ 0.01.

HQ exposure has previously been reported to elicit inflammatory responses in several cell types, potentially contributing to AMD-relevant pathology [27]. In our co-culture system, however, qPCR analysis did not detect significant changes in the expression of pro-inflammatory interleukins IL-6, IL-8, IL-1β in retinal samples. Similarly, Iba1 immunostaining did not reveal significant changes in the number or ONL migration of microglial cells (**Supplementary Figure 1**). These data suggest that the HQ-induced photoreceptor phenotype observed here is not accompanied by a robust inflammatory or microglial activation response detectable with these markers.

Altogether, the differential proteomic profile induced by ML240, along with the preservation of cytochrome c enrichment in photoreceptor inner segments, supports an adaptive neuroretinal response characterized by increased antioxidant-associated proteins, reduced ERAD/proteasome-associated stress markers, and improved mitochondrial integrity. This response was accompanied by preservation of photoreceptor structure in HQ-stressed co-cultures (**Figure 8**).

**Figure 8.**
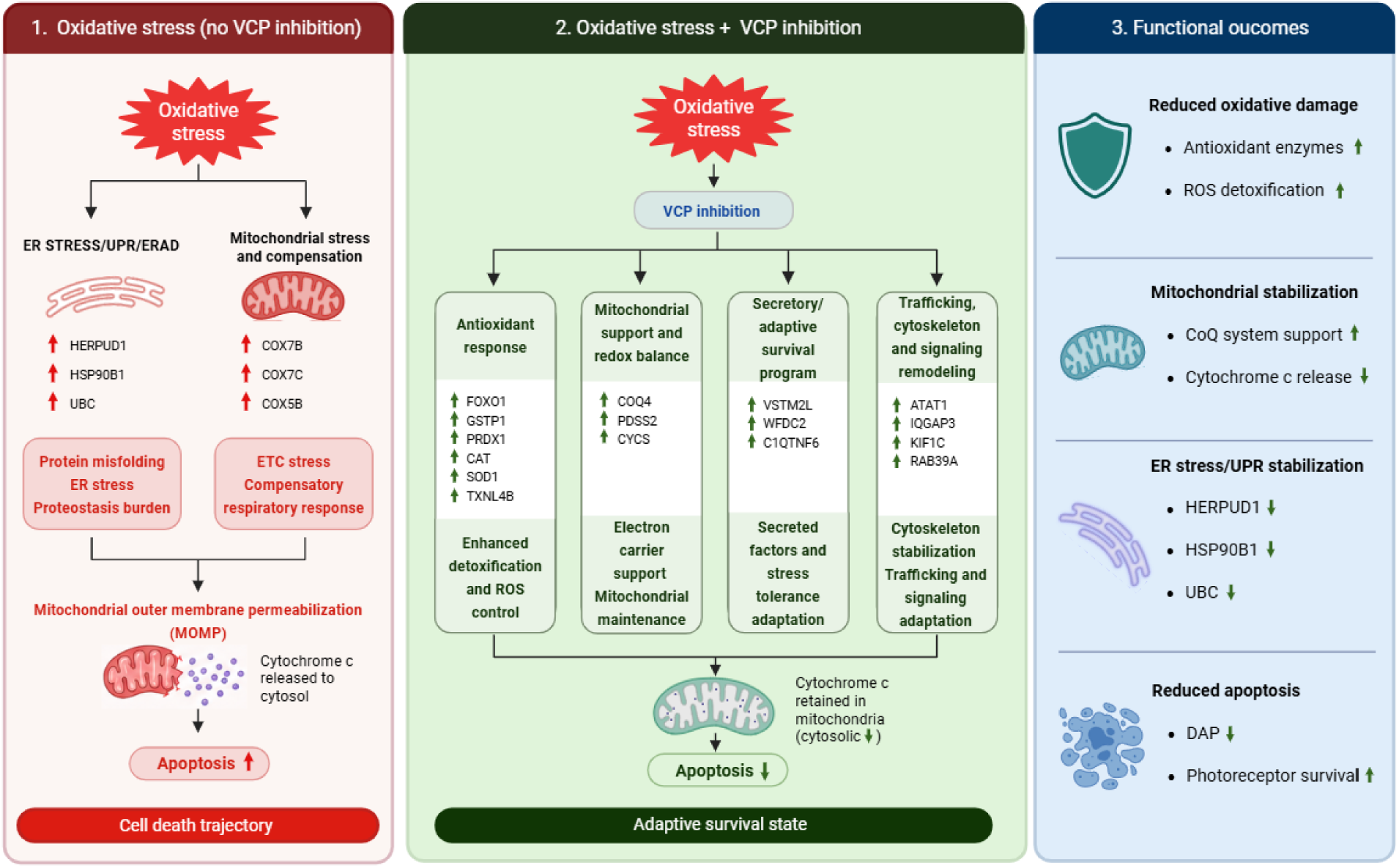
Graphical model of the neuroretinal response to ML240 in HQ-stressed co-cultures. Under HQ-induced stress, neuroretinas show increased abundance of ER stress/UPR-and ERAD-associated proteins, including HSP90B1, HERPUD1, and UBC, along with increased respiratory chain-associated proteins such as COX5B, COX7B, COX7C. HQ exposure is also associated with increased cytochrome c mislocalization from photoreceptor inner segments toward the ONL, consistent with compromised mitochondrial integrity, together with photoreceptor degeneration. By contrast, ML240 treatment shifts this response toward increased abundance of antioxidant-and redox-associated proteins (FOXO1, GSTP1, PRDX1, CAT, SOD1, TXNL4B, KEAP1), mitochondrial support-associated proteins (COQ4, PDSS2, CYCS), proteins linked to secretory or extracellular signaling responses (VSTM2L, WFDC2, C1QTNF6), and cytoskeletal or vesicle-trafficking proteins (ATAT1, IQGAP3, KIF1C, RAB39A). These changes are consistent with reduced proteostatic stress, enhanced antioxidant capacity, improved mitochondrial integrity, attenuated cytochrome c mislocalization, and preservation of photoreceptor structure under HQ-induced stress. Created with BioRender.

## Discussion

This study describes a human iPSC-RPE/porcine neuroretina co-culture model exposed to HQ, a lifestyle-associated stressor contributing to AMD. HQ induced caspase activation, oxidative stress, and apoptotic markers in iPSC-RPE cells, and caused photoreceptor degeneration in the co-cultured porcine neuroretina. Pharmacological inhibition of VCP did not attenuate the HQ-induced activation of apoptosis in iPSC-RPE cells, but selectively preserved photoreceptor structure, including OS length and cone density. This selective neuroretinal protection was associated with a shift toward antioxidant-, mitochondrial-, and cytoskeletal homeostatic protein profiles, together with a reduction of cytochrome c mislocalization in photoreceptors exposed to ML240.

RPE dysfunction is a root cause of AMD pathogenesis [28], yet rod photoreceptors are the first to die in advanced AMD [29]. HQ induces apoptosis, reduced ONL rows, decreased OS length, and lower cone density in the porcine neuroretina, features that capture neuroretinal changes in AMD. In particular, RPE stress and photoreceptor degeneration are critical hallmarks of AMD [30]. Nevertheless, this model should not be interpreted as a full reproduction of AMD, which is a chronic, multifactorial disease involving RPE, photoreceptors, Bruch’s membrane, choriocapillaris, immune pathways, and genetic susceptibility. Rather, the co-culture system presented in this study provides a reductionist tissue-level platform to investigate how environmental or systemic stressors affect the RPE-neuroretina axis as well as a high-content platform to test cell type-specific protective strategies.

A central finding of this study is the apparent cell type-specific effect of VCP inhibition. ML240 did not reduce HQ-induced caspase activity, oxidative stress, or cleaved caspase-3 induction in iPSC-RPE cells, yet it reduced stress response patterns in photoreceptors and preserved their structure in the co-cultured neuroretina. This suggests that RPE and photoreceptors differ in their stress response, their stress-adaptive capacity, or the therapeutic window in which VCP inhibition becomes protective rather than detrimental. These results also highlight the importance of using more complex models than monocultures, such as iPSC-RPE/neuroretina co-cultures, to dissect cell type-specific mechanisms of retinal stress and neuroprotection.

Proteomic analysis provided insight into this selective neuroretinal protection. HQ-treated neuroretinas showed increased abundance of proteins associated with ER stress, UPR, ERAD, and ubiquitin-proteasome pathways, including HERPUD1, HSP90B1, and UBC [31–33] compared with HQ plus ML240-treated neuroretinas. These changes are consistent with proteostatic stress in HQ-exposed retinal explants. Conversely, the lower abundance of these markers in ML240-treated explants suggests that VCP inhibition reshapes, rather than simply suppressing, the neuroretinal proteostasis response under oxidative stress. This is in line with previous studies showing that partial VCP inhibition can promote photoreceptor survival in retinal models characterized by proteostasis imbalance [21–23].

The proteomic data also indicate that ML240 modifies the neuroretinal response to oxidative stress. HQ-treated explants showed increased abundance of several mitochondrial respiratory chain-associated proteins, including COX7B, COX7C, and COX5B. Although this may reflect stress-associated mitochondrial remodeling [34], functional studies would be required to determine whether oxidative phosphorylation or ATP production are altered. In contrast, HQ plus ML240-treated retinas showed increased abundance of antioxidant and redox-associated proteins, including FOXO1, GSTP1, PRDX1, CAT, SOD1, and TXNL4B [35–40]. These changes suggest an enhanced antioxidant-associated protein profile that may contribute to photoreceptor resilience under HQ-induced stress.

The concomitant increase in KEAP1 is notable, given its role as a negative regulator of NRF2-dependent antioxidant responses [41]. One possible explanation for the upregulation of the antioxidant response despite the increase in KEAP1 could be that oxidative modification of KEAP1 alters its ability to restrain NRF2 by modifying its cysteine residues [42]. Furthermore, FOXO1-dependent pathways may also contribute to antioxidant gene regulation in a NRF2-independent manner. These hypotheses require direct validation in future studies.

In parallel, ML240-treated neuroretinas showed increased abundance of mitochondrial support-associated proteins, including COQ4 and CYCS [43, 44]. Immunostaining further showed that ML240 attenuated HQ-induced cytochrome c mislocalization from photoreceptor inner segments toward the ONL, while total TOM20 levels remained unchanged. These findings are consistent with improved mitochondrial integrity and reduced cytosolic release of cytochrome c, which is known to reduce apoptosis [45].

VCP acts as a central regulator of proteostasis, ER-associated degradation, organelle quality control, and mitochondrial homeostasis, processes that are particularly relevant in aging and retinal degeneration [46–48]. In this context, the increased abundance of proteins linked to mitochondrial homeostasis and cytoskeletal or vesicle-trafficking pathways, including ATAT1, IQGAP3, KIF1C, and RAB39A [49–51], suggests that the modulation of VCP activity may support structural and trafficking adaptations in stressed photoreceptors. Because photoreceptors are highly polarized neurons with intense proteostasis and energy demands, even modest shifts in protein quality control, mitochondrial stability, and intracellular transport may have a substantial impact on OS maintenance and photoreceptor preservation.

The lack of detectable induction of IL-6, IL-8, IL-1β, or Iba1-positive microglial migration suggests that the HQ-induced photoreceptor phenotype in this system was not accompanied by a robust inflammatory response. This contrasts with reports of HQ-driven inflammation in endothelial cells [27] but is compatible with the notion that effects of HQ are cell type- and context-dependent. Indeed, HQ has been reported to suppress inflammatory responses and macrophage activity also in other systems [52]. Broader cytokine profiling and microglial morphology or transcriptomic analyses would be required to assess inflammatory contributions in this model more comprehensively.

Several limitations should be acknowledged. As HQ was applied to the complete co-culture system, the present experiments do not distinguish between direct HQ toxicity to the neuroretina and secondary photoreceptor damage mediated by stressed iPSC-RPE cells. Second, although the human iPSC-RPE/porcine neuroretina co-culture preserves key tissue-level interactions that are absent from monocultures, it remains an *ex vivo* interspecies model and does not fully reproduce the chronic human *in vivo* environment of the retina, including choroidal vasculature, Bruch’s membrane remodelling, systemic immune inputs, lipid deposition and long-term aging-related changes. Finally, HQ exposure represents an acute oxidative insult rather than the chronic, multifactorial stress underlying AMD. Further studies assessing the effect of chronic stress in combination with an AMD-associated genetic background would help to elucidate long-term effects of stress in the retina and potential therapeutic strategies.

In conclusion, this study shows that HQ-induced oxidative stress causes iPSC-RPE damage and photoreceptor degeneration in a human iPSC-RPE/porcine neuroretina co-culture model, and that pharmacological VCP inhibition with ML240 selectively preserves photoreceptor structure under these conditions. The protective phenotype is associated with increased antioxidant and mitochondrial support-associated protein abundance, reduced ERAD/proteasome-associated stress markers, and attenuated cytochrome c mislocalization in photoreceptors, suggesting that the impact of VCP inhibition is both molecularly complex and cell-specific.

## Materials and methods

### iPSC-RPE generation and culture

The iPSC line iPS(IMR90)-4 (WiCell), heterozygous for CFH 402YH, was engineered to obtain the homozygous CFH 402YY genotype using CRISPR-Cas9 technology, as previously described [53–56]. Briefly, the single-guide RNA (sgRNA) sequence ATAGACGTTGCCTGCCATCC was synthesized using the *in vitro Guide-it™ sgRNA In Vitro Transcription and Screening Systems* (Takara). A total of 2 x 10^5^ iPSCs were electroporated with 2.2 µg Cas9 protein, and 450 ng sgRNA (5:1 mass ratio) in 10 µl of suspension buffer R using the Neon Transfection System (Invitrogen) [55], by two pulses of 1200 V and 20 ms. Cells were then plated onto laminin-coated 24-well plates in mTeSR1 media supplemented with 10 µM of ROCK inhibitor. After 48 h, cells were replated at low density (̴ 150 cells/cm^2^), to allow clonal colony formation. For the first 72 h, cells were fed with a 1:1 mixture of fresh mTeSR1 medium and conditioned medium collected from 60-80% confluent cells. Single colonies were collected manually with a p200 micropipette and replated into individual wells of laminin-coated 96-well plates for further expansion [55]. Single-cell-derived clones were screened by Sanger sequencing using the primers:

F: 5’GAAAATGTTATTTTCCTTATTTGGAAAATGG 3’and R: 5’GACACGGATGCATCTGGGA 3’. Positive clones with no indels were expanded to be differentiated into iPSC-RPE cells.

iPSCs were cultivated on hESC-matrigel-coated plates (354277, Corning) and seeded on hESC matrigel (356237, Corning) for differentiation into RPE cells according to a previously described protocol [13]. Briefly, iPSC were cultured in differentiation medium composed of DMEM/F12 (31331, Gibco), N2 supplement (17502, Gibco), B27 supplement (17504, Gibco), non-essential amino acids (NEAA; 11140, Gibco), and KnockOut Serum Replacement (KnockOut SR; 10828, Gibco). The medium was supplemented as follows: nicotinammide, 10 mM, days 0-4 (NO636-100G, Sigma-Aldrich), Noggin, 50 ng/ml, days 0-4 (1967-NG-025/CF, R&D systems), DKK-1, 10 ng/ml, days 0-6 (5439-DK-010/CF, R&D Systems), IGF-1, 10 ng/ml, days 0-6 (1291-G1-200, R&D Systems), bFGF, 5 ng/ml, days 2-4 (AF-100-18B, Peprotech), activin A, 100 ng/ml, days 4-14 (120-14E, Peprotech), SU5402, 10 μM, days 6-14 (sc-204308, Santa Cruz).

On day 14, cells were enriched by using TrypLE (12604; Thermo Fisher Scientific) according to manufactureŕs instructions in medium containing ROCK inhibitor Y27632 (StemMACS), and seeded onto Geltrex-coated (A1413302, Gibco) or hESC-qualified Matrigel-coated plates for expansion. Cells from passages 2 to 4 were seeded in the required experimental format, including 12-well culture inserts, 96-well plates, and 12-well plates, and matured for at least 50 days before experiments.

### Transepithelial electrical resistance (TER) measurement

TER was measured using a Millicell-ERS-2 voltohmmeter according to the manufacturer’s instructions. Electrodes were placed on the apical and basal sides of iPSC-RPE culture inserts, the current was applied, and resistance values (Ω) were recorded in triplicate for each insert. The final TER (Ω·cm²) was calculated by multiplying the measured resistance by the insert area.

### PEDF ELISA

Culture media from the apical and basal compartments of iPSC-RPE culture inserts was collected, and cellular debris was removed by centrifugation at 500 × g for 5 min. PEDF levels were quantified using a human PEDF ELISA kit (RD191114200R, BioVendor) following the manufacturer’s protocol. Standards, quality controls, and samples were incubated for 1 h at room temperature on a shaker. After washing, biotin-labelled antibody was added and incubated for 1h, followed by streptavidin-HRP conjugate for 1h. The reaction was developed with TMB substrate, stopped after 5 min, and absorbance was read at 405 nm using a Spark multimode microplate reader (Tecan, Switzerland).

### Porcine neuroretina isolation and co-culture

Adult porcine eyes were obtained from the local slaughterhouse and transported to the laboratory on ice. All procedures complied with European regulations for the use of animal by-products for research (Regulation (EC) No. 1069/2009; Regulation (EU) No. 142/2011) and were approved by the Landratsamt Tübingen under registration number DE 08 416 1157 21.

Eyes were dissected by removing the anterior segment and vitreous. The visual streak was identified based on the retinal vascular pattern, and 5mm biopsy punches were collected from this region. The neuroretina was carefully separated from the underlying RPE/choroid before transfer to culture inserts. Explants were transferred onto membrane inserts (353180, Corning-Falcon), placed in 12-well plates (353503, Corning-Falcon) containing pre-seeded mature iPSC-RPE monolayers. Explants were oriented with the photoreceptor side facing the iPSC-RPE monolayer.

Co-cultures were maintained for up to 8 days *in vitro* in serum-free medium (DMEM/F12:DMEM, 1.5:1) supplemented with 2 % B27, 1 % N2, and 1 % antibiotic–antimycotic (Gibco). Medium was changed every two days. Treatments were applied throughout the culture period and included 0.4% dimethyl sulfoxide (DMSO, vehicle control, A994.1, Roth), 20µM ML240 (5153, Tocris), 100µM hydroquinone (HQ, H9003, Sigma-Aldrich), or a combination of ML240 and HQ.

### *In vitro* cytotoxicity assays

Cytotoxicity and the activity of two key executioner caspases activated during apoptosis were evaluated using the MultiTox-Fluor Multiplex Cytotoxicity Assay (G9201, Promega) and the Caspase-Glo® 3/7 Assay (G8091, Promega), respectively, according to manufactureŕs instructions. Membrane damage was quantified based on cleavage of the bis-AAF-R110 (bis-alanylalanyl-phenylalanyl-rhodamine 110) dye, producing a fluorescent signal (Ex 485 nm/Em 520 nm). After fluorescence measurement, Caspase-Glo reagent was added, and luminescence was recorded using a Spark multimode microplate reader (Tecan, Switzerland).

Reactive oxygen species (ROS)-associated hydrogen peroxide production was assessed using the ROS-Glo H₂O₂ Assay (G8820, Promega). Cells were incubated with the H₂O₂ substrate for 4 h, followed by addition of the detection buffer containing cysteine and signal enhancer. Luminescence was measured after 10 min using a Spark multimode microplate reader.

### Histology

Co-cultured explants were fixed in 4% paraformaldehyde (PFA) in 0.1M phosphate buffer (PB, pH 7.4) for 45 minutes and cryoprotected in a sucrose gradient (10%, 20%, and 30%). Samples were embedded in cryomatrix (Tissue-Tek® O.C.T. Compound, 4583, Sakura® Finetek, VWR) and snap-frozen in liquid nitrogen. Radial sections (14 µm) were cut, air-dried at 37°C, and stored at -20°C. For iPSC-RPE monocultures, cells were fixed in 4% PFA in PBS for 45 min, washed three times with PBS for 5 minutes, and stored in PBS at 4°C.

### Immunohistochemistry

iPSC-RPE monocultures or 14 µm co-cultured retina/RPE sections were pre-treated with TrueBlack solution (23007, Biotium), diluted 1:80 in 70% ethanol, to reduce autofluorescence. Samples were washed three times for 5 min with PBS, permeabilized, and blocked for 1h at room temperature in 0.1% PBST, PBS containing 0.1% Tween-20, supplemented with 10% normal goat serum (311-053, PAA-Labs) or donkey serum (D9663, Sigma), and 1% bovine serum albumin (K41-001, PAA-Labs). Samples were incubated overnight at 4°C with the primary antibodies diluted in blocking solution (anti-ZO-1 1:100, 610966, BD Biosciences; anti-cleaved caspase 3 1:400, 9661, Cell Signalling; anti-Rhodopsin 1:350, MAB5316, Sigma Aldrich; anti-M opsin 1:200, AB5405, Sigma Aldrich; anti-cytochrome c 1:500, ab13575, Abcam; anti-iba1 1:500, 019-19741, Fujifilm Wako Chemicals). Appropriate species-specific secondary antibodies conjugated to Alexa Fluor 488 or Alexa Fluor 568, (Alexa Fluor™568 dye-conjugated goat anti-mouse IgG, 1:500, A11031, ThermoFisher; Alexa Fluor™ 568 dye-conjugated goat anti-rabbit IgG, 1:500, A11036, Molecular Probes) were applied for 1 h at room temperature, followed by DAPI counterstaining. Samples were mounted with Fluoromount-G (17984-25, Electron Microscopy Sciences).

### Microscopy and image analysis

Z-stack images were acquired using a Zeiss Axio Imager Z1 ApoTome microscope (20x or 63x objective) and analysed with Zen Blue 3.7 software. Three to four sections per sample were imaged. ONL rows were manually counted based on DAPI staining. OS length was measured based on rhodopsin staining in five evenly distributed positions per image and averaged. Cone density was calculated as cones per 100 µm of retinal section, averaging three different regions per image. Cytochrome C mean fluorescence intensity (MFI) was measured across the ONL and OS/IS layers, IS/ONL MFI ratio was calculated. Microglia activation was assessed by Iba-1 immunostaining, quantifying the number of Iba1-positive cells located within or migrating toward the ONL. All image post-processing adjustments (contrast, brightness) were applied equally across groups.

### Mass Spectrometry (MS)

Retinal explants were isolated from the IPSC-RPE cells and collected in lysing kit tubes (P000933-LYSK0-A, Precellys) with 100 µl of lysing buffer (87787, Thermo Fisher Scientific) to which we added protease and phosphatase inhibitor (1861281, Thermo Fischer Scientific). Lysates were homogenised using a Precellys 24 tissue homogenizer. After 20 minutes of centrifugation at 4°C, supernatants were collected and protein concentration was quantified by Bradford (500-0006, Biorad). 20 µg of lysates were taken and an acetone-based protein precipitation was performed followed by tryptic digest and desalting of digested proteins via stop-and-go extraction tips (Thermo Fisher Scientific) as previously described [57] and prepared for mass spectrometry analysis.

LC-MS/MS analyses were performed at the Core Facility for Medical Proteomics, University of Tübingen, on a nanoElute 2 nano-flow LC system coupled to a timsTOF Ultra 2 mass-spectrometer (Bruker Daltonics). For each sample, 1 µL of peptide samples was loaded onto a Bruker PepSep C18 column (25 cm × 75 µm i.d., 1.5 µm particle size) and separated at a flow rate of 300 nL/min, with the column maintained at 50 °C in a column oven. Peptides were eluted using mobile phase A (0.1% formic acid in water) and mobile phase B (0.1% formic acid in acetonitrile) with the following gradient: 5% B at 0 min, linearly increased to 25% B over 24 min, to 35% B at 30 min, and to 95% B at 34 min; 95% B was held until 38 min, after which the column was re-equilibrated to 5% B by 40 min, giving a total run time of 40 min.

Mass spectra were acquired in dia-PASEF mode [58] using a CaptiveSpray ion source (Bruker Daltonics) operated at 1600 V, with an MS1 scan range of 100–1700 m/z. Twenty-four dia-PASEF windows (12 × 2) were defined for serial MS2 fragmentation, covering an m/z range of 327.5– 1400 and an ion mobility range of 0.60–1.45 1/K₀; collision energies were ramped linearly from 20 to 59 eV across this ion mobility range.

Raw MS data were analyzed with DIA-NN v1.8.1 [59] in library-free mode against the human UniProt database. A precursor ion library was first generated from an in silico FASTA digest combined with deep learning–based spectral prediction. The experimental library produced by the initial DIA-NN search was then used for cross-run normalization and mass accuracy correction. Protein quantification was restricted to high-confidence identifications, applying a precursor false discovery rate (FDR) threshold of 0.01 and considering only tryptic peptides with up to two missed cleavages. The match-between-runs option was enabled, and shared spectra were excluded from protein identification.

### Western Blot

Retinal lysates were obtained and quantified as described above. A total of 20 µg protein per sample was mixed with 1x Laemmli sample buffer, separated on 8%–16% SDS-PAGE gels (XP08165Box; Thermo Fisher Scientific) and transferred onto PVDF membranes (0,45µm, 10600023, Cytiva). Membranes were blocked in 5% milk in TBST (Tris-buffered saline with 0.1% Tween 20) and incubated with the corresponding primary antibodies (anti-cyt c 1:1000, 556433, BD Biosciences; anti-TOM20 1:1000, 42406T, Cell Signalling; anti-β-actin 1:2000, 3700S, Cell Signalling) overnight at 4°C. Membranes were incubated for 1 h at room temperature with horseradish peroxidase-coupled (HRP)-conjugated secondary antibodies (anti-mouse 1:2000, 7076P2, Cell Signalling; anti-rabbit 1:2000, 7074P2, Cell Signalling). ECL Plus chemiluminescent HRP detection reagent (32132, Thermo Fisher Scientific) was applied, and signal was detected using a FusionFX imaging system (Vilber Lourmat, France). Band intensities were quantified using ImageJ software and normalized to β-actin, and relative protein expression levels were determined based on the control group.

### RT-qPCR

Neuroretinal explants were isolated from the iPSC-RPE cells, and RNA was extracted using PureZOL (732-6880, Bio-Rad). Samples were homogenized by inversion and incubated at room temperature for 5 min. Chloroform was added, and samples were vortexed for 15 s, incubated for 5 min at room temperature, and centrifuged at 12,000 g for 15 min at 4°C. The aqueous phase was collected and mixed with isopropanol for RNA precipitation. Samples were centrifuged at 12,000 g for 15 min at 4°C. Pellets were rinsed twice with 75% EtOH, air-dried, and resuspended in 20 μl of RNase-free water. RNA purity and concentration were measured using Nanodrop spectrophotometer. cDNA was synthesized via reverse-transcription of 0.5 μg of RNA using SuperScript™ II Reverse Transcriptase (18064-071, Thermo Fisher Scientific), random primers (Y02321, Thermo Fisher Scientific), and dNTPs (U151B, Promega) in a total volume of 23 µl following the manufactureŕs instructions. Differential gene expression was analysed by qRT-PCR using SYBR Green Master Mix (4309155, Thermo Fisher Scientific) and gene-specific forward and reverse primers (IL-6 fwd 5’-CGGTCTTGTGGAGTTTCAGATA -3’, rev 5’-CTGGATCAGTGCTTTGGTACT -3’; IL-1β fwd 5’-TCCAGGACAAAGACCACAAATC -3’, rev 5’-GCAGAACACCACTTCTCTCTTC -3’; IL-8 fwd 5’-GAAGCAACAACAACAGCAGTAA -3’, rev 5’-GCACAGGAATGAGGCATAGA -3’; β-actin fwd 5’-AAGCCAACCGTGAGAAGATG -3’, rev 5’-GCAGAAGAAAGAACAAGTGAGAAAG -3’).

### Statistical analyses

Immunohistochemistry, Western blot, and RT-qPCR data were analyzed using GraphPad Prism version 10. Data are presented as mean ± SEM. Each data point represents one biological replicate (n). For each explant, multiple radial sections and non-overlapping regions were imaged and quantified. Technical measurements from each explant were averaged to generate one value per biological replicate. Normality was assessed using the Shapiro-Wilk test. Comparisons between two groups were performed using paired or unpaired Student’s t-tests, according to the experimental design. For experiments involving more than two groups, one-way ANOVA followed by Tukey’s multiple comparisons test was used. A p-value < 0.05 was considered statistically significant.

Downstream statistical analysis for MS data was performed in Perseus v1.6.15.0 [60]. Label-free quantification (LFQ) intensities were filtered for valid values in at least 3 replicates in at least one group. LFQ intensities from HQ and HQ-ML240 were normalized to the median of the DMSO group, batch corrected using the ComBat method implemented in Perseus and compared between groups using a two-sided permutation-based t-test (250 permutations, p < 0.05, with multiple-testing correction) to identify putative differentially abundant proteins. For pathway enrichment analysis, only proteins exceeding a log₂ fold-change threshold of 0.38 were considered differentially regulated. Enrichment against the Reactome database was performed using Enrichr [61], and all downstream visualizations were generated using custom Python scripts.

## Acknowledgments

The authors would like to thank Dr. Sven Schnichels and his team from the Institute of Ophthalmic Research in Tübingen for facilitating porcine material.

## Author contribution statement

All authors contributed substantially to the work of this paper. ACAG, BAG, and MU designed the study. BAG and MU supervised the study. ACAG, AA, SA, EC, RFG, BC, ASPD, AV, and SB performed the experiments. ACAG, AA, SA, and MAJ analysed and interpreted the data. ACAG drafted the manuscript, and AA, EK, BAG, and MU reviewed the manuscript. Corresponding author: BAG

## Ethic statement

All procedures involving porcine tissue were performed in accordance with institutional guidelines and were approved by the local competent authority (Landratsamt Tübingen), under the registration number DE 08 416 1157 21, authorizing the use of category 1 animal by-products for research purposes (Regulation (EC) No. 1069/2009; Regulation (EU) No. 142/2011).

## Data Availability statement

The raw data generated in this study have been deposited to the ProteomeXchange Consortium via the MassIVE partner repository (MSV000102782, doi:10.25345/C5K35MV0C) with the dataset identifier PXD082362.

