## Supplementary material for "VCP inhibition preserves photoreceptor integrity under hydroquinone-induced oxidative stress in a human iPSC-RPE/porcine neuroretina co-culture model": Almansa-Garcia et al 2026b Supplementary material.docx

S1. Inflammatory markers in the co-cultured porcine retina

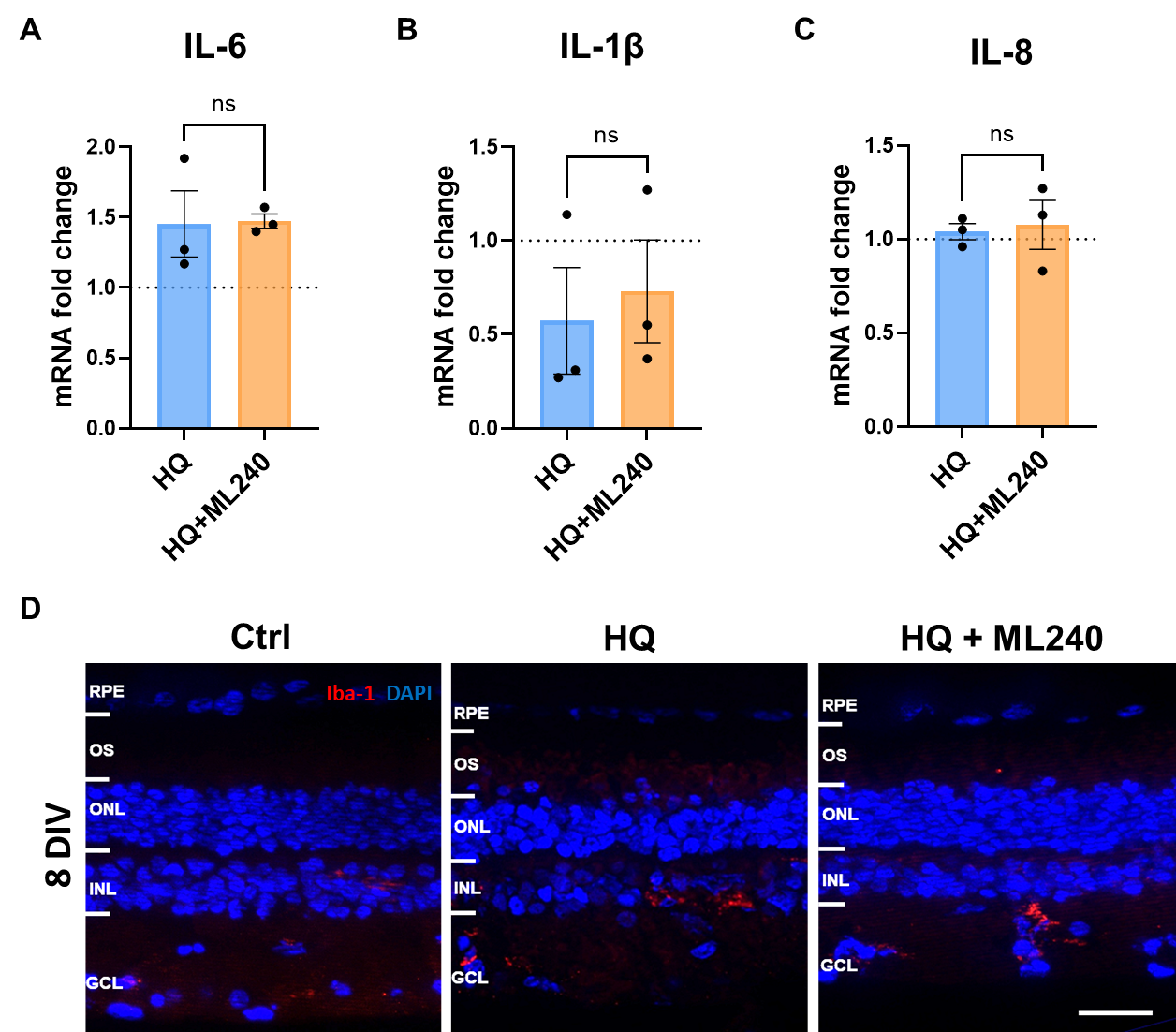

**Supplementary figure 1. HQ does not induce robust inflammatory marker changes in co-cultured retinas.** mRNA fold change assessed by qPCR for **(A)** interleukin 6 (IL-6), (**B**) IL-1β, and (**C**) IL-8. (**D**) Immunofluorescence staining of Iba1-positive microglia (red) in co-cultured retinas treated with HQ, HQ + ML240, or DMSO vehicle control for 8 days (DIV 8). Data presented as mean ± SEM. The control mean is indicated by a dotted line for reference. n = 3. ns = no significance. Scale bar 20 µm.

**S2. Full-length WBs**

**Figure 2D:**

**BCL2**

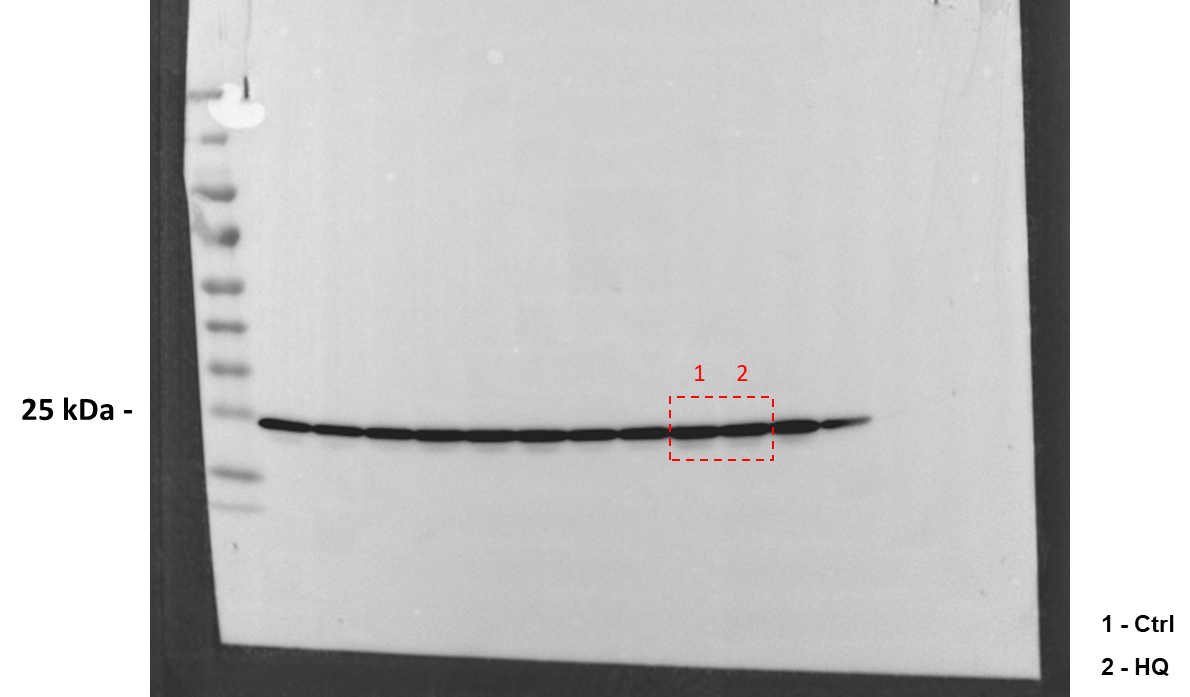

**BAX**

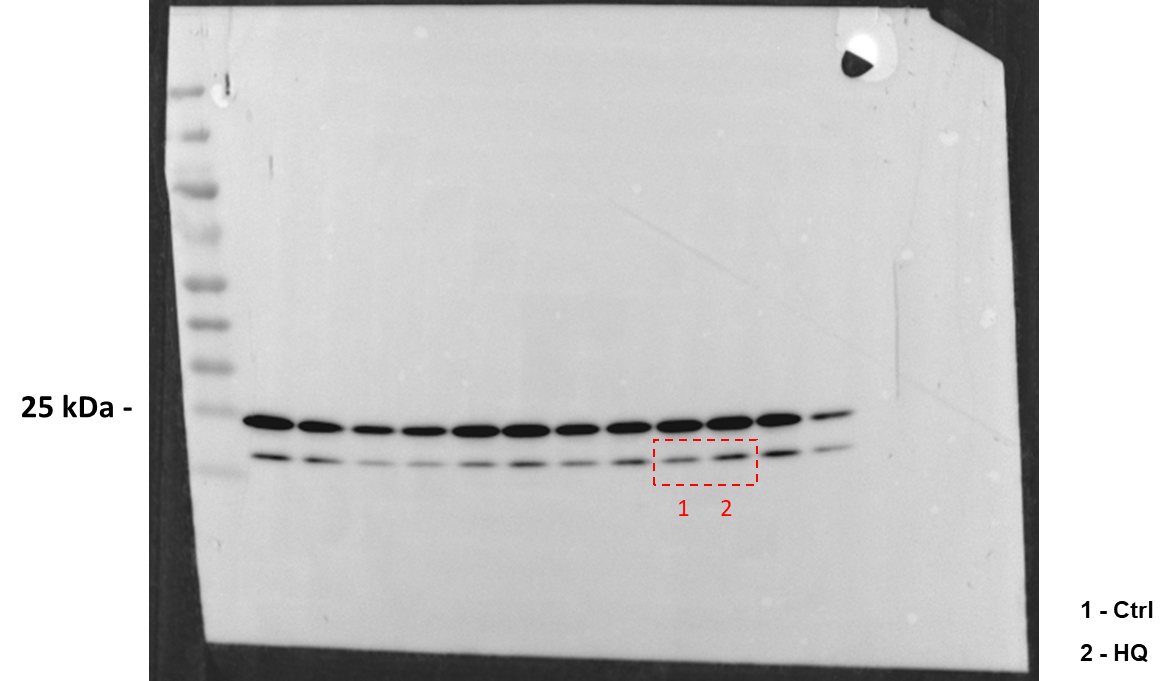

**Actin of BCL2 and BAX**

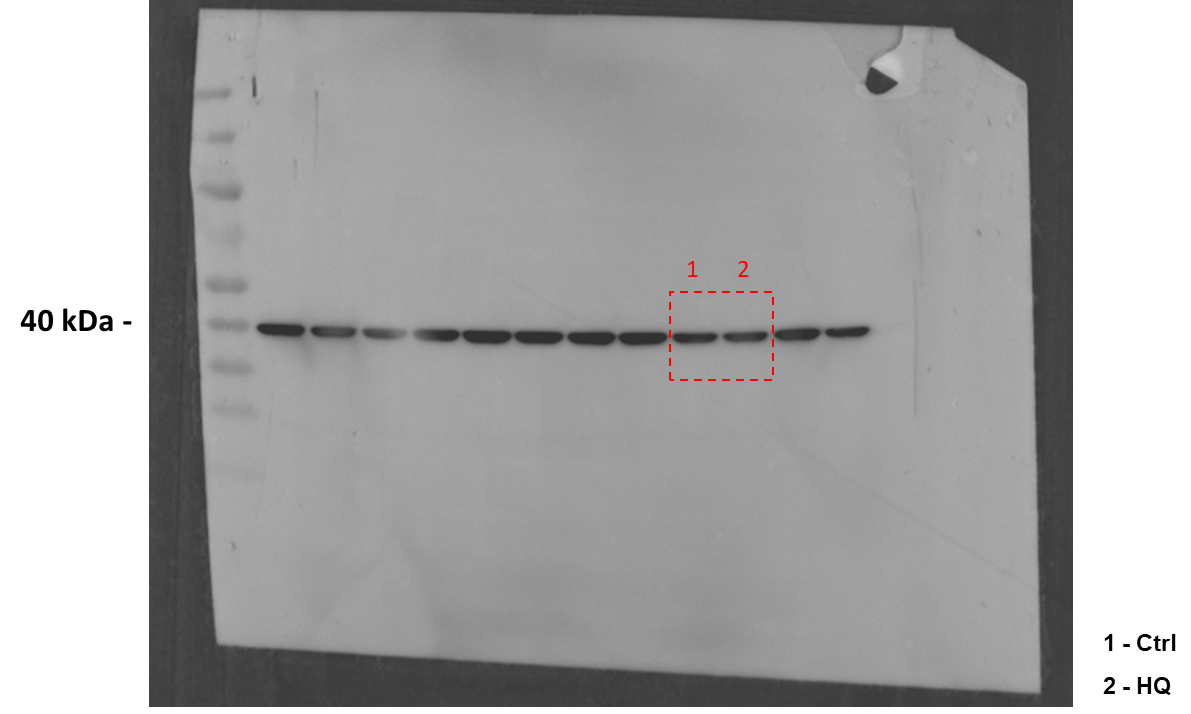

**Figure 7A:**

**Cyt c**

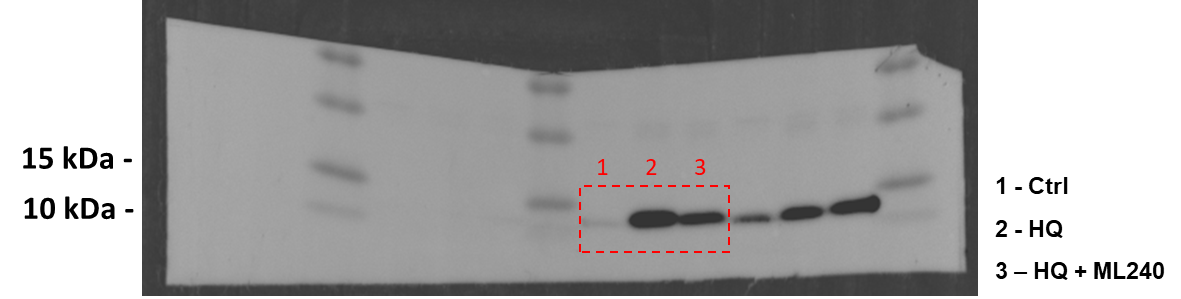

**TOM20**

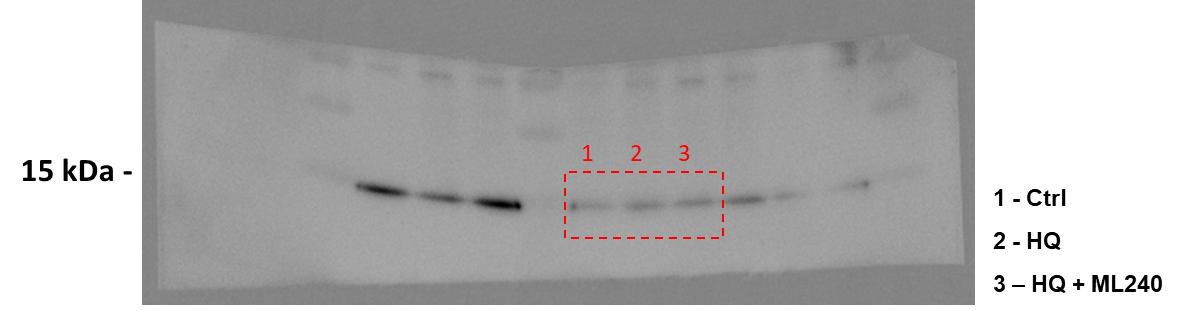

**Actin of Cyt C and TOM20**

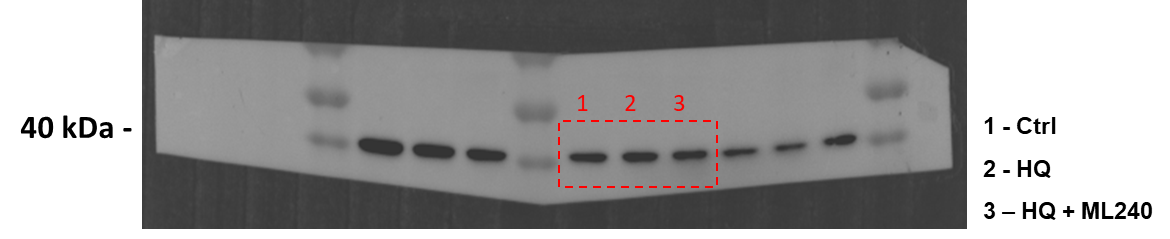
